# Miltefosine enhances infectivity of miltefosine-resistant *Leishmania infantum* by attenuating innate immune recognition

**DOI:** 10.1101/2021.02.22.432409

**Authors:** Dimitri Bulté, Lieselotte Van Bockstal, Laura Dirkx, Magali Van den Kerkhof, Carl De Trez, Jean-Pierre Timmermans, Sarah Hendrickx, Louis Maes, Guy Caljon

## Abstract

Miltefosine (MIL) is currently the only oral drug available to treat visceral leishmaniasis but its use as first-line monotherapy has been compromised by an increasing treatment failure. Despite the scarce number of resistant clinical isolates, MIL-resistance by mutations in a single aminophospholipid transporter gene can easily be selected in a laboratory environment. These mutations result in a reduced survival in the mammalian host, which can partially be restored by exposure to MIL, suggesting a kind of drug-dependency. To enable a combined study of the infection dynamics and underlying immunological events for differential *in vivo* survival, firefly luciferase (PpyRE9) / red fluorescent protein (DsRed) double-reporter strains were generated of MIL-resistant (MIL-R) and syngeneic MIL-sensitive (MIL-S) *Leishmania infantum*. Results in C57Bl/6 and BALB/c mice show that MIL-R parasites induce an increased innate immune response that is characterized by enhanced influx and infection of neutrophils, monocytes and dendritic cells in the liver and elevated serum IFN-γ levels, finally resulting in a less efficient establishment in liver macrophages. The elevated IFN-γ levels were shown to originate from an increased response of hepatic NK and NKT cells to the MIL-R parasites. In addition, we demonstrated that MIL could increase the *in vivo* fitness of MIL-R parasites by lowering NK and NKT cell activation, leading to a reduced IFN-γ production. These data provide an immunological basis for the MIL-R-associated attenuated phenotype and for the peculiar drug-dependency that may constitute one of the mechanisms of treatment failure.

**Importance:** Recently, our laboratory demonstrated an *in vivo* fitness loss of experimentally selected MIL-R parasites in both the sand fly vector and vertebrate host. These findings could explain the scarce number of MIL-R clinical isolates. Surprisingly, MIL-R parasites developed a MIL-dependency which could partially rescue their fitness loss and which may constitute a mechanism of treatment failure. This research aimed to better understand the immunological basis of the attenuated phenotype and the effect of MIL on infectivity traits. Together, this study provides new insights into the complex interplay between the parasite, drug and host and discloses an immune-related mechanism of treatment failure.

## Introduction

Visceral leishmaniasis (VL) is a fatal vector-borne disease that is caused by transmission of the protozoan parasite *Leishmania donovani* in Asia and parts of Africa and by *L. infantum* in the Mediterranean area and Latin America (1–3). In 2017, 90 000 new cases of VL were reported with another 300 million people at risk of acquiring the disease (4–6). *Leishmania* parasites are transmitted through the bites of infected female sand flies. During blood feeding, extracellular promastigotes are injected into the dermis of the host where they are taken up by cells of the phagocytic system. Inside these cells, promastigotes differentiate into obligate intracellular amastigotes that will disseminate and infect macrophages in the major target organs liver and spleen (7–9). In most VL mouse models, different outcomes of infection can be distinguished depending on the tissue (10). In the liver, an acute infection leads to the development of granulomas around infected Kupffer cells (resident macrophages of the liver) resolving the infection. The cytokine interferon (IFN)-γ plays a crucial role in this process by stimulating the production of tumor necrosis factor (TNF)-α and inducible nitric oxide synthase (iNOS) in infected Kupffer cells (11). On the other hand, the immune response in the spleen is characterized by high TNF-α levels and a delayed or absent granuloma formation leading to a chronic infection accompanied by architectural damage and immunological dysfunction (10,12–18). Early immunological events play a crucial role in shaping these organ-specific immune responses. In the liver, one of those events is the recruitment and activation of natural killer (NK) and NKT cells within hours of infection, as seen during experimental infections with *L. donovani* and *L. infantum* (19–22). Whereas IFN-γ-producing CD4^+^ and CD8^+^ T cells are crucial for parasite clearance at a later stage of the infection, IFN-γ production by NK and NKT cells during the early onset is critical for efficient granuloma formation and early parasite control (19, 22). While several reports have shown that NK cells are not essential in shaping the early immune response, the absence of NKT cells or NKT cell activation result in decreased IFN-γ levels, impaired granulomatous response and higher parasite burdens in both the liver and spleen (21, 22).

A successful vaccine against VL has not been developed to date and drug treatment has been hampered by widespread resistance, treatment failure (TF) and toxicity of the current antileishmanial drugs (23). Since its introduction in 2002, miltefosine (MIL) has remained the only oral drug available for VL treatment (23–25). Although the exact mode of action of MIL is not entirely understood, changes in the parasite’s lipid metabolism, reduction of mitochondrial cytochrome c oxidase and induction of apoptosis-like cell death are at the basis of its activity (26–31). Rather soon after its introduction, relapse rates increased up to 20% (32, 33) with indication of an elevated drug-tolerance (34) but without the emergence of full and stable MIL-resistance. To date, only few fully MIL-resistant (MIL-R) isolates have been described, including two MIL-R *L. infantum* strains isolated from French HIV-infected patients and two MIL-R *L. donovani* strains recovered from patients in India (35–37). Despite the scarce number of MIL-R field isolates (33–39), resistance can easily be selected experimentally arising from mutations in the *MIL transporter* (*MT*) or the *Ros3* gene (37, 40). The encoded MT/Ros3 transporter complex facilitates MIL import over the parasite’s cell membrane (41). Mutations in either the *MT* or *ROS3* gene have been reported in the four above mentioned MIL-R field isolates (35–37). Re-introduction of functional *MT* or *ROS3* gene copies fully restores MIL-susceptibility (37), linking the defective subunits to the MIL-R phenotype. Former research by our lab and others has indicated that *MT* mutations provoke a severe fitness loss of *Leishmania* strains in mice and sand flies (40,42–45). This fitness loss was characterized by a reduced *in vitro* growth rate and metacyclogenesis and a reduced IL-10 production by infected macrophages (43). Furthermore, MIL-R parasites were unable to cause a typical visceralization pattern in mice, with rapid clearance from the viscera (42). In the sand fly vector, these parasites showed a reduced colonization of the stomodeal valve by metacyclic promastigotes, a process that is essential for transmission to the vertebrate host (45). These observations were recently confirmed by our lab using a natural MIL-R *L. infantum* strain (LEM5159) which has a defective *LiROS3* gene caused by a frameshift mutation and early stop codon (46). Surprisingly, the reduced infectivity in mice was partially restored either following *in vivo* MIL treatment or by drug pre-exposure of parasites (42). To gain better insights into the impact of resistance acquisition and mechanisms underlying apparent MIL-dependency of MIL-R parasites, the present study evaluated the immunological characteristics of early infections with and without MIL-exposure.

## Results

### MIL-R parasites fail to disseminate during the first week of the infection

Compared to susceptible (MIL-S) parasites, isogenic MIL-R parasites were rapidly cleared from the liver in C57Bl/6 and BALB/c mice with a failure to successfully disseminate to the spleen and bone marrow (Fig. 1). MIL-S*^PpyRE9/DsRed^* showed a typical visceralization pattern with a rapid increase of hepatic parasite burdens during the first weeks of the infection followed by dissemination of parasites to the spleen and bone marrow. In contrast, hepatic MIL-R*^PpyRE9/DsRed^* burdens were the highest at day 1 of infection after which they rapidly subsided. Spleen and bone marrow signals were only rarely detected during onset of MIL-R*^PpyRE9/DsRed^* infection. Flow cytometry (Fig. 2A) revealed a higher number of infected cells in the liver at one day post-infection (dpi) of MIL-R*^PpyRE9/DsRed^* versus MIL-S*^PpyRE9/DsRed^* infected C57Bl/6 mice (p≤0.05). Within 3 weeks of infection, these numbers steadily declined in MIL-R*^PpyRE9/DsRed^* but stagnated in MIL-S*^PpyRE9/DsRed^* infected mice. In the spleen, the total number of infected cells was approximately 5-fold lower than in the liver with no major differences between MIL-R*^PpyRE9/DsRed^* and MIL-S*^PpyRE9/DsRed^*. Non-DsRed MIL-S*^PpyRE9^* and MIL-R*^PpyRE9^* controls were included at each time point to validate whether the observed DsRed signals indeed corresponded with infected cells rather than infection-induced changes in autofluorescence (Fig. S1). It must be noted that the number of infected cells (Fig. 2A) as well as the intensity of the DsRed signal (Fig. S1) decreases over the course of infection. While this was expected for the MIL-R*^PpyRE9/DsRed^*, it was not for the MIL-S*^PpyRE9/DsRed^* strain given that the bioluminescence data clearly indicated a parasite expansion. This increase of MIL-S*^PpyRE9/DsRed^* parasite burdens was confirmed by Giemsa counting of liver and spleen tissue imprints at 1, 7 and 21 dpi (Fig. 2B). The lower DsRed reporter signals are therefore most likely associated with the increasingly quiescent state of intracellular amastigotes.

**Figure 1:**
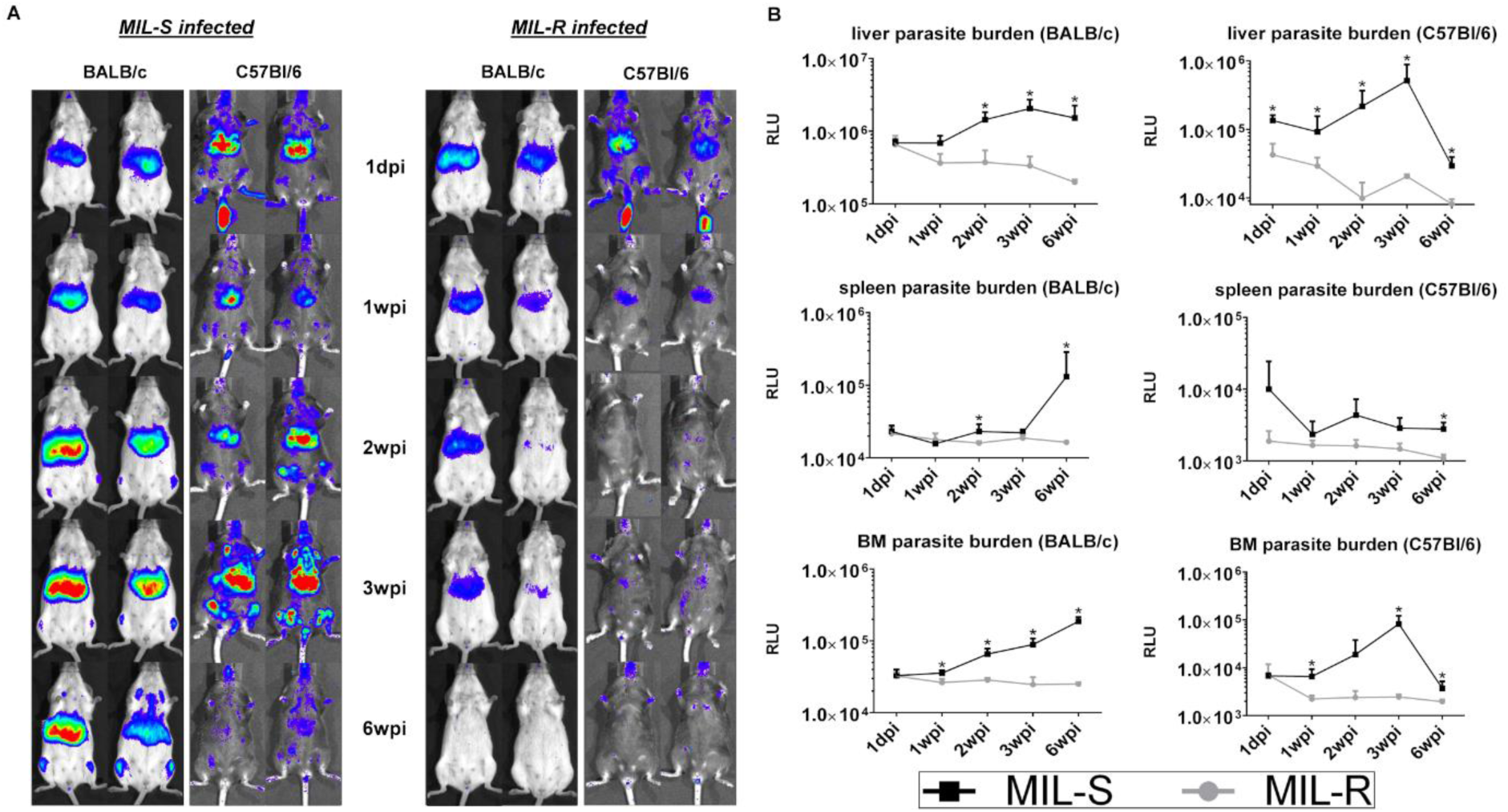
Infection dynamics of MIL-S and MIL-R in BALB/c and C57Bl/6 mice. **(A)** Bioluminescence imaging, using an exposure time of 15 minutes, of MIL-S*^PpyRE9/DsRed^* (left panel) and MIL-R*^PpyRE9/DsRed^* (right panel) infected BALB/c and C57Bl/6 mice during the first 6 weeks of infection showing a reduced infectivity of MIL-R*^PpyRE9/DsRed^* parasites. **(B)** Mean RLU values of liver, spleen and bone marrow (BM) bioluminescent signal during the first 6 weeks of MIL-S*^PpyRE9/DsRed^* and MIL-R*^PpyRE9/DsRed^* infection in BALB/C (left) and C57Bl/6 (right) mice. Each infection group consisted of 3 BALB/c or C57Bl/6 mice and at least 3 independent repeats were performed. Results are expressed as mean ± SD (* p≤0.05).

**Figure 2:**
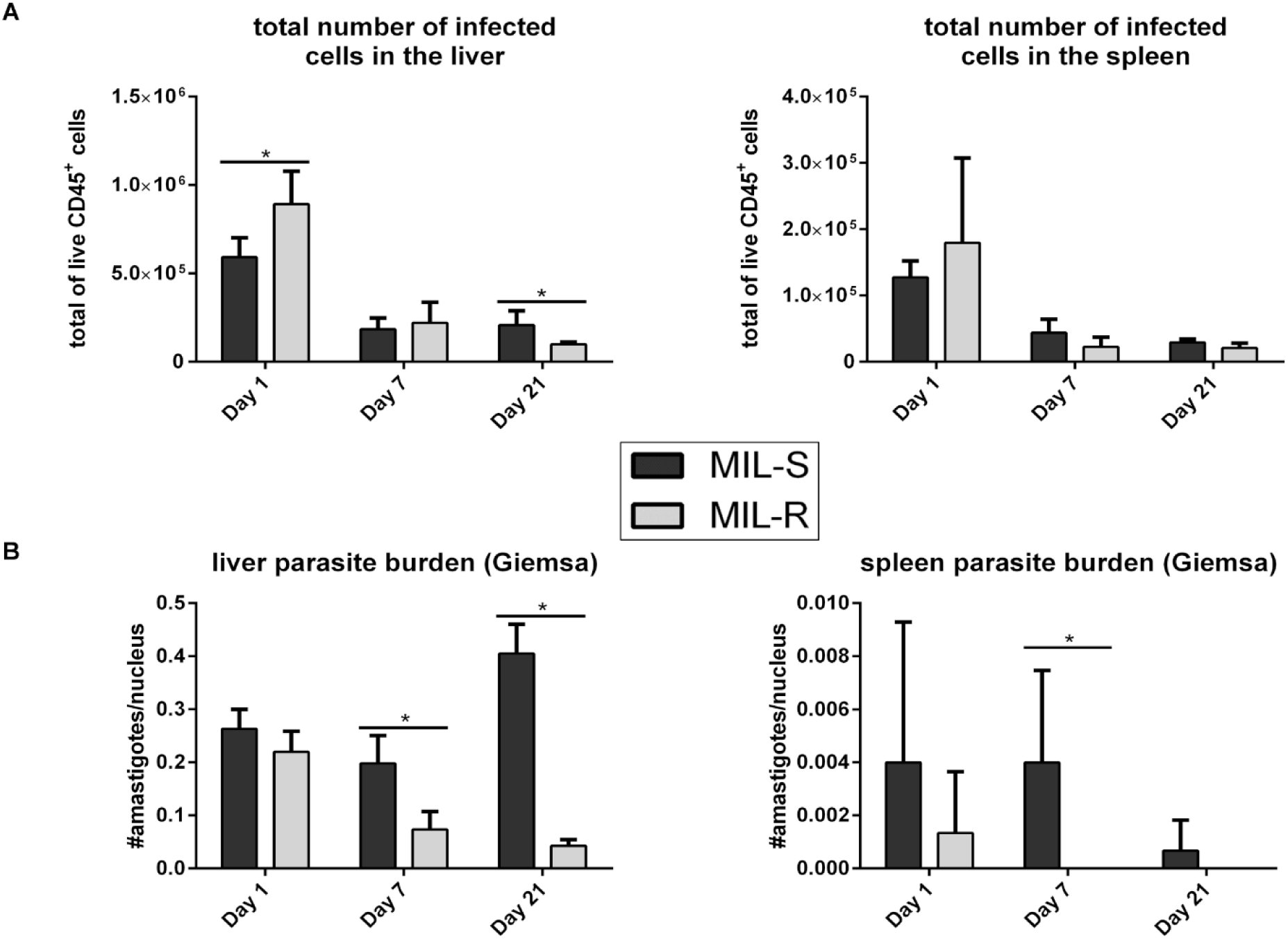
Parasite burdens in the liver and spleen following MIL-S and MIL-R infection in C57Bl/6 mice. (A) Total number infected cells in the liver and spleen of C57Bl/6 mice infected with MIL-S*^PpyRE9/DsRed^* and MIL-R*^PpyRE9/DsRed^* at 1, 7 and 21 dpi. (B) At 1, 7 and 21 dpi, tissue imprints of MIL-S*^PpyRE9/DsRed^* and MIL-R*^PpyRE9/DsRed^* infected livers and spleens were Giemsa stained and the number of amastigotes per nucleus was determined by microscopic counting. Experiments were carried out in triplicate with 3 to 5 mice per group and three independent repeats were performed. Results are expressed as mean ± SD (* p≤0.05).

### Myeloid cells are rapidly recruited to the liver and spleen during early MIL-R infection resulting in an altered cellular distribution

An increased influx of monocytes, neutrophils and dendritic cells was observed in MIL-R*^PpyRE9/DsRed^* infected livers at 1 dpi (Fig. 3A) with a different cellular distribution of infection (Fig. 3B). In MIL-R*^PpyRE9/DsRed^* infected livers, monocytes (CD11b^+^ Ly-6G^-^ Ly-6C^High^ F4/80^-^), neutrophils (CD11b^+^ Ly-6G^+^) and dendritic cells (CD11b^+^ CD11c^+^) represented a higher percentage of the total number of DsRed^+^ CD11b^+^ cells when compared to the MIL-S*^PpyRE9/DsRed^* (28.0 ± 10.6% vs 16.3 ± 3.9%; 6.6 ± 3.4% vs 3.1 ± 0.7%; 8.3 ± 1.7% vs 3.6 ± 0.9%; p ≤0.05) while the relative importance of macrophages (CD11b^+^ Ly-6G^-^ Ly-6C^low^ F4/80^+^) diminished (43.7 ± 11.2% vs 68.7 ± 6.7%; p≤0.05). At 7 and 21 dpi, the differences in cellular distribution between MIL-S*^PpyRE9/DsRed^* and MIL-R*^PpyRE9/DsRed^* became less prominent; however, monocytes, neutrophils and dendritic cells still represented a higher percentage of the total DsRed^+^ CD11b^+^ cells in MIL-R*^PpyRE9/DsRed^* infected liver. In the spleen no major differences in myeloid cell influx and cellular tropism were observed (Fig. S2A-B). These data show that MIL-R*^PpyRE9/DsRed^* induces a stronger influx of myeloid cells in the liver during the initial phase of infection with concomitant change of initial cell tropism.

**Figure 3:**
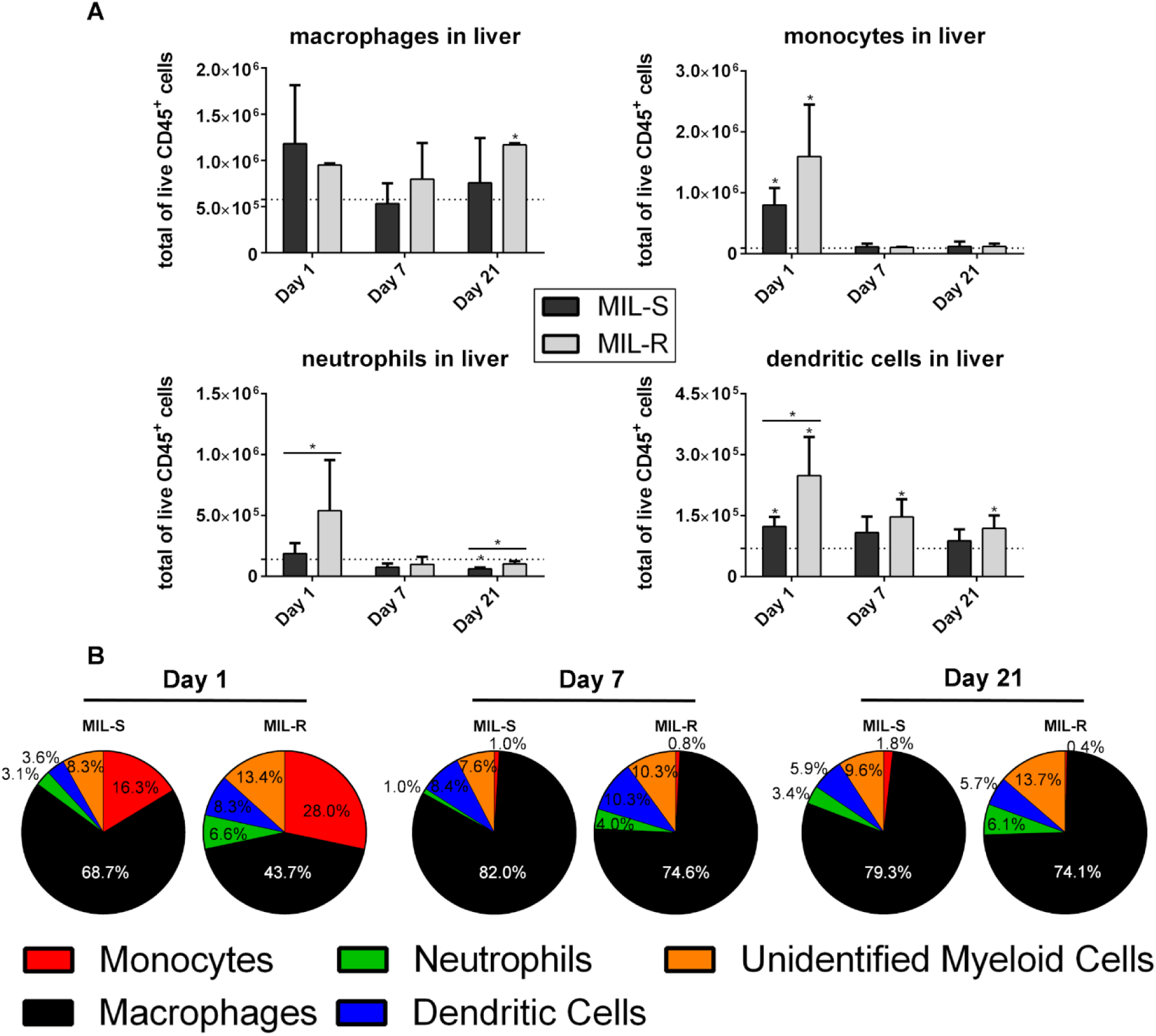
Flow cytometry analysis of the myeloid compartment of the liver. (A) Total number of macrophages, monocytes, neutrophils and dendritic cells in the liver of C57Bl/6 mice infected with MIL-S*^PpyRE9/DsRed^* and MIL-R*^PpyRE9/DsRed^* at 1, 7 and 21 dpi. Dotted lines indicate the average number of cells in naive animals. (B) Distribution of infected cells among the CD11b^+^ myeloid compartment in the livers of MIL-S*^PpyRE9/DsRed^* and MIL-R*^PpyRE9/DsRed^* infected C57Bl/6 mice at 1, 7 and 21 dpi. Experiments were carried out in triplicate with 3 to 5 mice per infection group and three independent repeats were performed. Results are expressed as mean ± SD (* p≤0.05).

At 7 and 21 dpi, different infected cell populations in the liver (macrophages, monocytes, neutrophils and dendritic cells) were sorted and analyzed microscopically for differences in the intracellular parasite burdens between MIL-S*^PpyRE9/DsRed^* and MIL-R*^PpyRE9/DsRed^* (Fig. 4). By 21 dpi, liver macrophages were more heavily infected by MIL-S*^PpyRE9/DsRed^* parasites compared to MIL-R*^PpyRE9/DsRed^* (5.52 ± 3.18 amastigotes/cell vs 3.08 ± 2.34 amastigotes/cell; p≤0.0001) whereas no differences could be observed in the other myeloid cell types.

**Figure 4:**
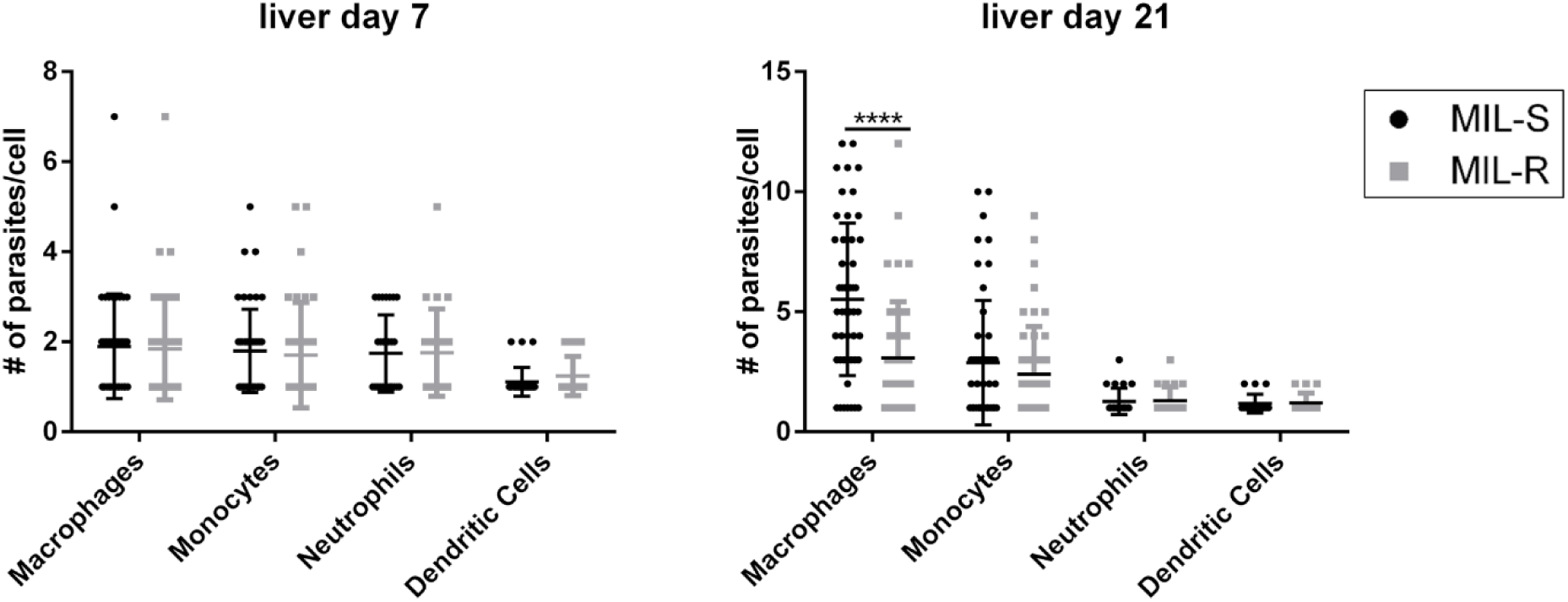
Infection ratios of sorted cell populations in the liver and spleen of MIL-S and MIL-R infected mice. Infected macrophages, monocytes, neutrophils and dendritic cells in the liver were sorted from C57Bl/6 mice and infection ratios were determined microscopically. Results are expressed as mean ± SD (**** p≤0.0001).

### MIL-R parasites induce stronger early IFN-γ responses in hepatic NK and NKT cells

At three time points during early infection (1, 7 and 21 dpi), serum levels of several infection-induced key inflammatory cytokines were determined (Fig. 5A). No major differences in TNF-α, IL-10 or IL-1β levels were observed. Both parasite strains induced a moderate TNF-α response at 1 dpi (43.5 ± 32.1 pg/mL and 51. 8 ± 29.6 pg/mL, respectively) and TNF-α remained elevated throughout the early infection with a second peak at 21 dpi (62.6 ± 12.8 pg/mL and 45.0 ± 26.9 pg/mL, respectively). IL-10 levels were moderately elevated throughout infection with both strains, with an initial increase at 1 dpi (29.4 ± 14.0 pg/mL and 37.7 ± 27.8 pg/mL for MIL-S*^PpyRE9/DsRed^* and MIL-R*^PpyRE9/DsRed^*, respectively) and a stronger increase at 3 wpi in MIL-S*^PpyRE9/DsRed^* compared to MIL-R*^PpyRE9/DsRed^* (59.8 ± 11.3 pg/mL and 27.8 ± 13.6 pg/mL, respectively). For IL-1β, levels did not exceed 10 pg/mL and no important interstrain differences were observed. As for IFN-γ, a prominent response (*p* <0.01) was observed following MIL-R*^PpyRE9/DsRed^* infection at 1 dpi (213.1 ± 123.1 pg/mL) in comparison to the moderate MIL-S*^PpyRE9/DsRed^*-induced IFN-γ response (62.0 ± 53.3 pg/mL). In both infection groups, the early IFN-γ surge subsided by 4 dpi. These results were confirmed by quantitative RT-qPCR of the gene expression levels of *IFN-γ*, *TNF-α*, *IL-10* and *IL-1β* in the liver and spleen (Fig. S3A). Enhanced transcription of the *IFN-γ* gene was observed at each time point for MIL-R*^PpyRE9/DsRed^* compared to MIL-S*^PpyRE9/DsRed^* in the liver and spleen, with statistical differences in the liver at 1 and 7 dpi. *TNF-α* transcription was significantly increased in MIL-R*^PpyRE9/DsRed^* infected livers at 1, 7 and 21 dpi and at 1 and 7 dpi in the spleen. Less interstrain variation was observed for the *IL-10* and *IL-1β* transcript levels.

**Figure 5:**
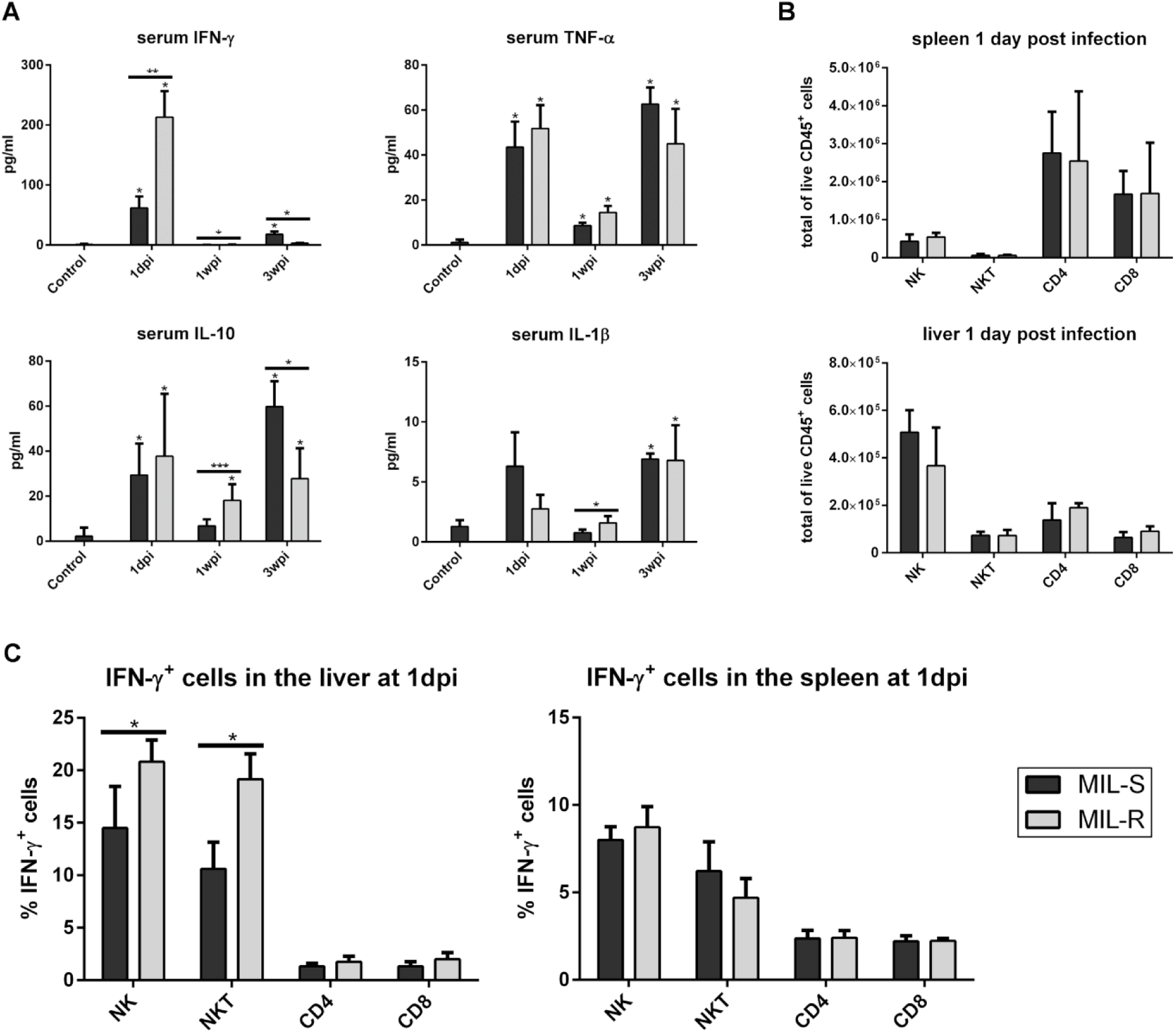
Serum cytokine levels during infection with MIL-S and MIL-R parasites. (A) Serum levels of IFN-γ, TNF-α, IL-10 and IL-1β at different time points during infection with MIL-S*^PpyRE9/DsRed^* and MIL-R*^PpyRE9/DsRed^* in C57Bl/6 mice. Combined results of 3 independent experiments with 3 to 5 mice per infection group. Results are expressed as mean ± SD. Significance levels relative to the non-infected controls are indicated above the bars. (B) Total number of NK, NKT, CD4^+^ T and CD8^+^ T cells in the liver (lower panel) and spleen (top panel) at 1 dpi. Experiments were carried out in triplicate with 3 to 5 mice per infection group. (C) Percentage of IFN-γ producing NK, NKT, CD4^+^ T and CD8^+^ T cells in the liver and spleen of uninfected, MIL-S*^PpyRE9/DsRed^* and MIL-R*^PpyRE9/DsRed^* infected IFN-γ reporter mice. Experiments were carried out in duplicate with 3 mice in each infection group. Results are expressed as mean ± SD. Significance levels relative to the non-infected controls are indicated above the bars (* p≤0.05, ** p≤0.01 and *** p≤0.001).

The increased IFN-γ levels seen at 1 dpi suggest an enhanced infiltration and/or activation of IFN-γ producing lymphocytes upon MIL-R*^PpyRE9/DsRed^* infection. It has been shown that different lymphocyte subsets produce IFN-γ throughout a VL infection in mice (48). While CD4^+^ and CD8^+^ T cells are an important source of IFN-γ during the later stages of infection when fully mature granulomas develop to clear infected Kupffer cells, NK and NKT cells were shown to produce IFN-γ during the initial stages of infection (19–22). Given the early IFN-γ response at day 1 of infection, NK (TcRβ^-^ NK1.1^+^), NKT (TcRβ^+^ CD1d tetramer^+^), CD4^+^ T (TcRβ^+^ CD4^+^) and CD8^+^ T (TcRβ^+^ CD8^+^) cell populations were examined in the liver and spleen of MIL-S*^PpyRE9/DsRed^* and MIL-R*^PpyRE9/DsRed^* infected C57Bl/6 mice. Both strains induced infiltration of similar numbers of NK, NKT, CD4^+^ T and CD8^+^ T cells at 1 dpi (Fig. 5B). IFN-γ reporter (GREAT) mice (49, 50) were used next to investigate the activation of these cells and to identify the source of differential IFN-γ production at 1 dpi, (Fig. 5C and S3B). Compared to an uninfected reporter control, both strains induced IFN-γ production by NK and NKT cells in the liver whereby MIL-R*^PpyRE9/DsRed^* induced more IFN-γ-producing NK and NKT cells. No differential induction of the IFN-γ reporter was observed in splenic NK and NKT cells at 1 dpi, nor was there any induction of IFN-γ in CD4^+^ and CD8^+^ T cells in the liver and spleen at this time point (Fig. 5C). These data clearly indicate that MIL-R parasites induce an enhanced activation of hepatic NK and NKT cells leading to an increased IFN-γ production at the very early onset of infection.

### NK and NKT cells contribute to early MIL-R parasite clearance

To investigate whether the early IFN-γ burst and increased NKT cell activation contributes to the decreased survival of MIL-R*^PpyRE9/DsRed^*, infections were carried out in BALB/c mice and/or C57Bl/6 treated with an anti-IFN-γ or an anti-CD1d antibody to neutralize CD1d-mediated lipid antigen presentation to NKT cells. BALB/c mice treated with an anti-IFN-γ neutralizing antibody 1 day before MIL-R*^PpyRE9/DsRed^* infection showed increased liver burdens as compared to isotype treated mice (Fig. 6). No effect was observed on spleen and bone marrow burdens. The contribution of NKT cells to the increased IFN-γ production during the first 24h of infection was confirmed by a drastic drop in IFN-γ serum levels as a result of anti-CD1d treatment (Fig. 7C). Bioluminescent imaging of the MIL-R*^PpyRE9/DsRed^* infection dynamics revealed significantly increased liver burdens in anti-CD1d as compared to isotype treated BALB/c and C57Bl/6 mice (Fig. 7A-B). In most anti-CD1d-treated mice, some dissemination to the spleen and bone marrow was observed. Similar results, but less pronounced, were obtained in anti-NK1.1-depleted C57Bl/6 mice showing an increased survival of MIL-R*^PpyRE9/DsRed^* during the first weeks of infection compared to isotype control-treated mice (Fig. S4). These data indicate the importance of early NK and NKT cell responses and the corresponding IFN-γ production in the rapid clearance of MIL-R parasites from the liver preventing further dissemination to other target organs. It must be noted that IFN-γ and NKT cell neutralization and NK cell depletion does not fully restore parasite survival, indicating that still other factors contribute to the decreased *in vivo* infectivity.

**Figure 6:**
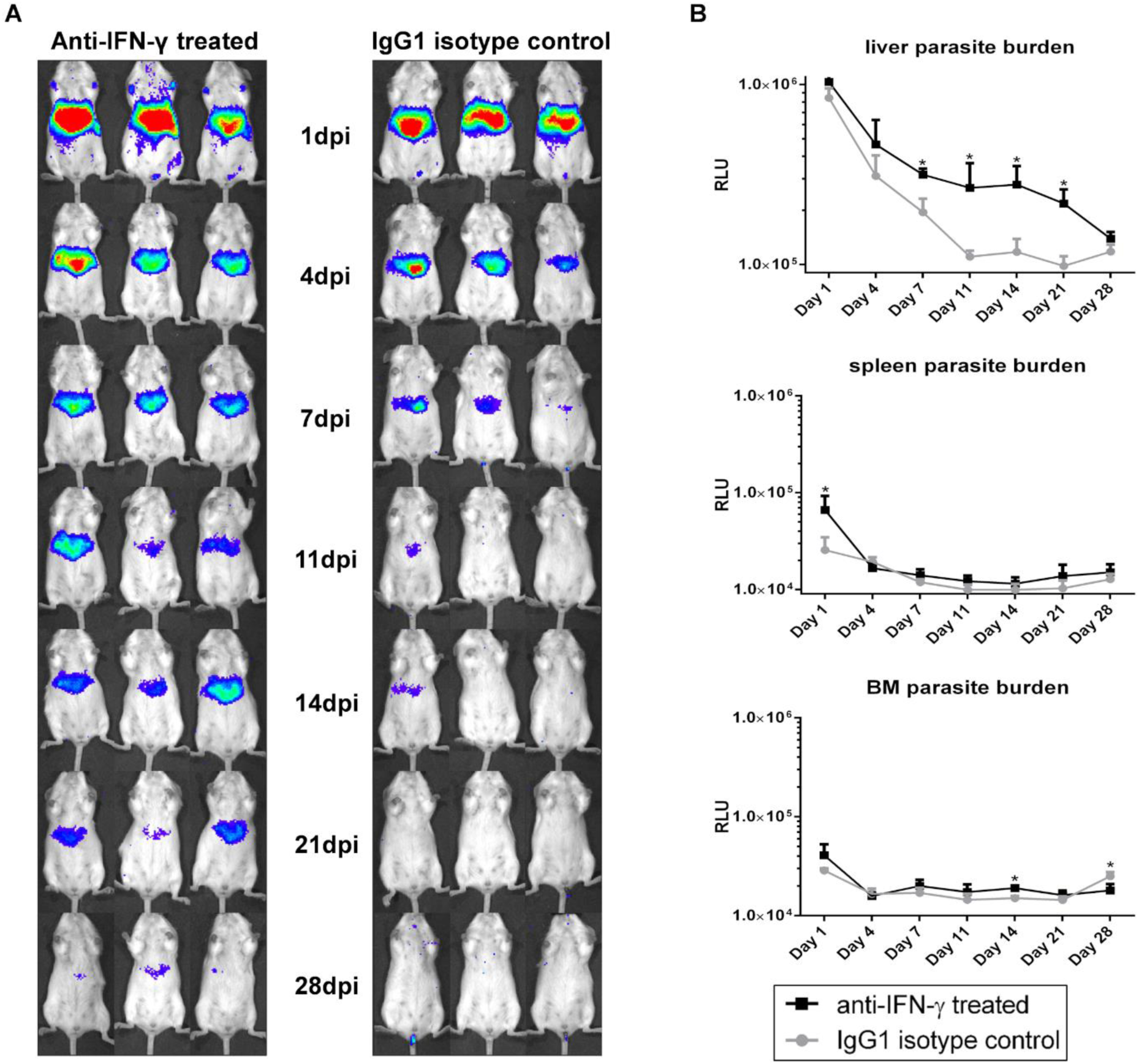
Effect of IFN-γ neutralization on early infection dynamics of MIL-R. (A) Bioluminescent imaging using an exposure time of 15 min of MIL-R*^PpyRE9/DsRed^* infected BALB/c mice treated with an anti-IFN-γ neutralizing antibody (left panel) or an IgG1 isotype control antibody (right panel) 1 day prior to infection. (B) Mean RLU values of liver, spleen and BM bioluminescent signals during the first 4 weeks of MIL-R*^PpyRE9/DsRed^* infection in anti-IFN-γ- and IgG1 isotype-treated BALB/C. Each infection group consisted of 3 BALB/c mice. Results are expressed as mean ± SD (* p≤0.05).

**Figure 7:**
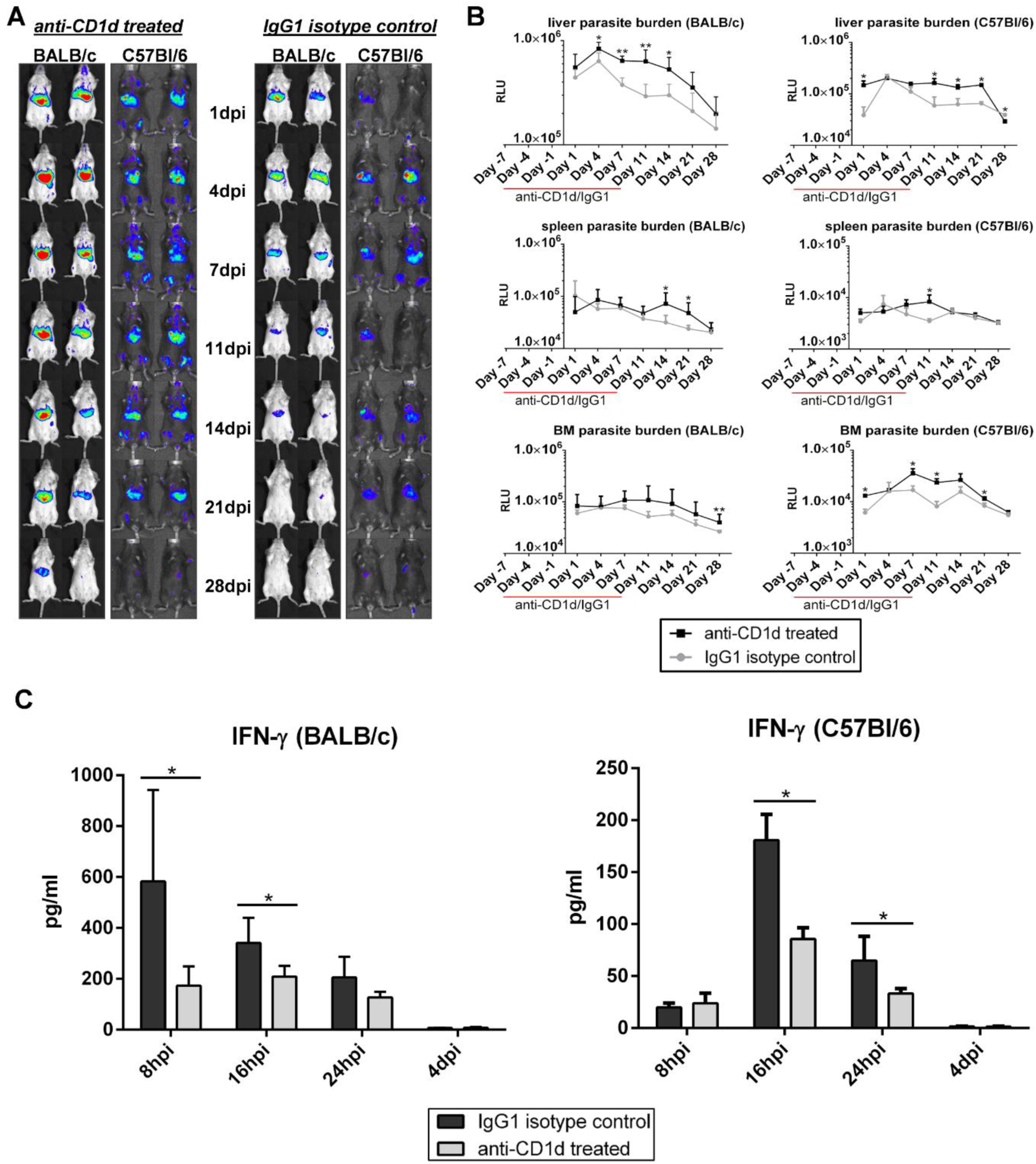
Early infection dynamics of MIL-R with or without CD1d neutralization. (A) Bioluminescent imaging using an exposure time of 15 min of MIL-R*^PpyRE9/DsRed^* infected BALB/c and C57Bl/6 mice treated with an anti-CD1d neutralizing antibody (left panel) or an IgG1 isotype control antibody (right panel) for 2 weeks starting 1 week before infection. (B) Mean RLU values of liver, spleen and BM bioluminescent signals during the first 4 weeks of MIL-R*^PpyRE9/DsRed^* infection in anti-CD1d- and IgG1 isotype-treated BALB/C and C57Bl/6 mice. (C) Comparison between the serum IFN-γ levels during the first 24h of MIL-R*^PpyRE9/DsRed^* infection in anti-CD1d- and IgG1 isotype-treated BALB/c and C57Bl/6 mice. For the experiments in BALB/c mice, each infection group consisted of 6 BALB/c mice and for the experiments in C57Bl/6 mice, two independent repeat experiments were performed with 3 mice in each infection group. Results are expressed as mean ± SD (* p≤0.05 and ** p≤0.01).

### Neutrophils have a dual role during the MIL-R infection onset

Given the infection-induced neutrophil influx, depletion was performed in BALB/c mice to study the impact on MIL-R*^PpyRE9/DsRed^* infection and early IFN-γ burst (Fig 8). MIL-R*^PpyRE9/DsRed^* parasites failed to establish following transient neutrophil depletion. Dissemination to the bone marrow and spleen was seen in the IgG2a isotype-treated mice but not in the neutrophil-depleted mice, indicating that neutrophils support MIL-R*^PpyRE9/DsRed^* dissemination. Furthermore, IFN-γ levels were significantly decreased in the absence of neutrophils during the first 24h of the infection compared to the levels in IgG2a-treated control mice. These data indicate that MIL-R parasites show a more reduced survival in neutropenic mice, in line with a protective role of neutrophils as initial host cell while contributing to triggering the initial IFN-γ response.

**Figure 8:**
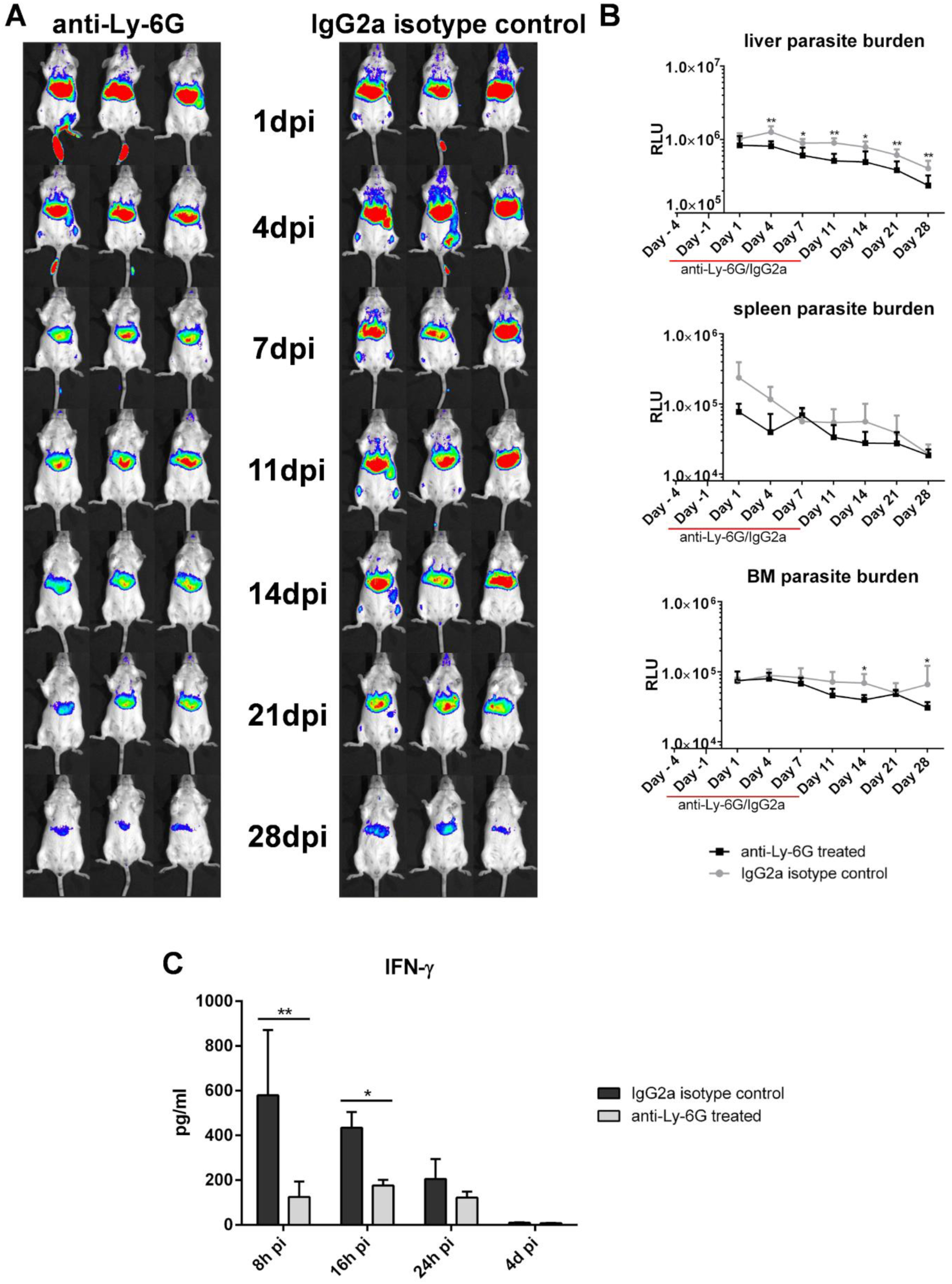
Early infection dynamics of MIL-R^PpyRE9/DsRed^ in neutropenic mice. (A) Bioluminescent imaging using an exposure time of 15 min of MIL-R*^PpyRE9/DsRed^* infected BALB/c mice treated with an anti-Ly-6G depleting antibody (left panel) or an IgG2a isotype control antibody (right panel) for 2 weeks starting 1 week before infection. (B) Mean RLU values of the liver, spleen and BM bioluminescent signals during the first 4 weeks of MIL-R*^PpyRE9/DsRed^* infection in anti-Ly-6G and IgG2a isotype treated BALB/c mice. (C) Comparison between the serum IFN-γ levels during the first 24h of MIL-R*^PpyRE9/DsRed^* infection in anti-Ly-6G and IgG2a isotype treated BALB/c mice. Each infection group consisted of 6 BALB/c mice. Results are expressed as mean ± SD (* p ≤0.05, ** p ≤0.01).

### MIL increases MIL-R parasite survival by reducing the activation of NK and NKT cells

We recently described that the MIL-R parasite developed a drug dependency that translates into an increased *in vivo* survival in the presence of MIL, either by *in vivo* treatment or *in vitro* pre-incubation of the parasite (42). Using bioluminescent imaging, increased survival was observed during the first weeks of infection with MIL-R*^PpyRE9/DsRed^* pre-incubated with MIL compared to a control MIL-R*^PpyRE9/DsRed^* infected group (Fig. 9A-B). After 2 weeks, parasite numbers rapidly dropped in both groups until no differences could be observed by week 4 of infection. In the previous sections, it was shown that MIL-R*^PpyRE9/DsRed^* induced a strong activation of NK and NKT cells leading to an increased IFN-γ response. The pre-incubation of MIL-R*^PpyRE9/DsRed^* with MIL resulted in a 6-fold reduction of serum IFN-γ levels at 1 dpi (135.6 ± 40.0 pg/ml vs 21.4 ± 12.0 pg/ml; Fig. 9C). This reduced IFN-γ production was found to be associated with a reduced activation of liver NK (p=0.1) and NKT cells (p< 0.05; Fig. 9D and Fig.S5). No induction of the IFN-γ reporter was observed in splenic NK and NKT cells at 1 dpi, nor was there any induction of IFN-γ in CD4^+^ and CD8^+^ T cells in the liver and spleen.

**Figure 9:**
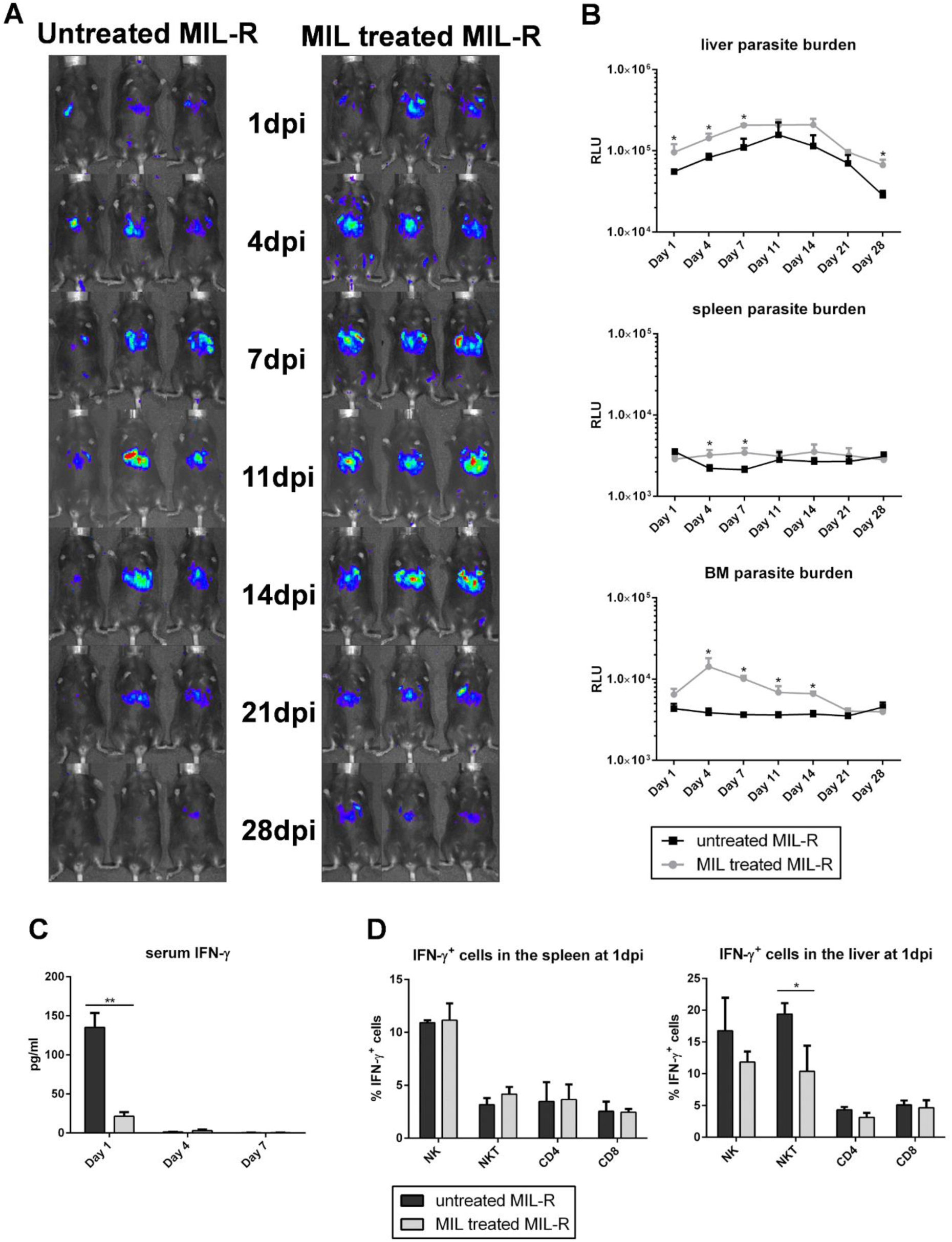
Impact of MIL pre-treatment on the MIL-R infectivity in mice. (A) Bioluminescent imaging using an exposure time of 15 min of MIL pre-treated MIL-R*^PpyRE9/DsRed^* and control MIL-R*^PpyRE9/DsRed^* infected C57Bl/6 mice. (B) Mean RLU values of the liver, spleen and BM bioluminescent signals during the first 4 weeks of MIL pre-treated MIL-R*^PpyRE9/DsRed^* and control MIL-R*^PpyRE9/DsRed^* infection in C57Bl/6 mice. Two independent repeat experiments were performed with 3 to 5 mice in each infection group. (C) IFN-γ serum levels during the first week of infection with MIL-pretreated MIL-R*^PpyRE9/DsRed^* and control MIL-R*^PpyRE9/DsRed^* in C57Bl/6 mice. (D) Percentage of IFN-γ producing NK, NKT, CD4^+^ T and CD8^+^ T cells in the liver and spleen of uninfected, MIL-S*^PpyRE9/DsRed^* and MIL-R*^PpyRE9/DsRed^* infected IFN-γ reporter mice. Infection groups consisted of 3 to 6 C57Bl/6 mice and experiments were performed in duplicate. Results are expressed as mean ± SD (* p ≤0.05, ** p ≤0.01).

### MIL increases the infectivity of MIL-R parasites in the skin after intradermal inoculation

To further confirm MIL-dependency of MIL-R parasites in more natural settings, the effect of MIL treatment was evaluated in BALB/c mice following intradermal (i.d.) inoculation with a natural MIL-R (LEM5159*^PpyRE9^*) parasite, harboring a mutation in the LiROS3 subunit of the MT transporter. This i.d. infection model is a better representation of the natural infection route, although parasites in this model fail to disseminate to the visceral organs. Therefore, the effect of MIL treatment was studied at the inoculum site in the ear (Fig. 10). *In vivo* MIL treatment of LEM5159*^PpyRE9^* infected mice resulted in higher parasite burdens at all time points during the first 4 weeks of the infection indicating MIL dependency of this natural MIL-R strain.

**Figure 10:**
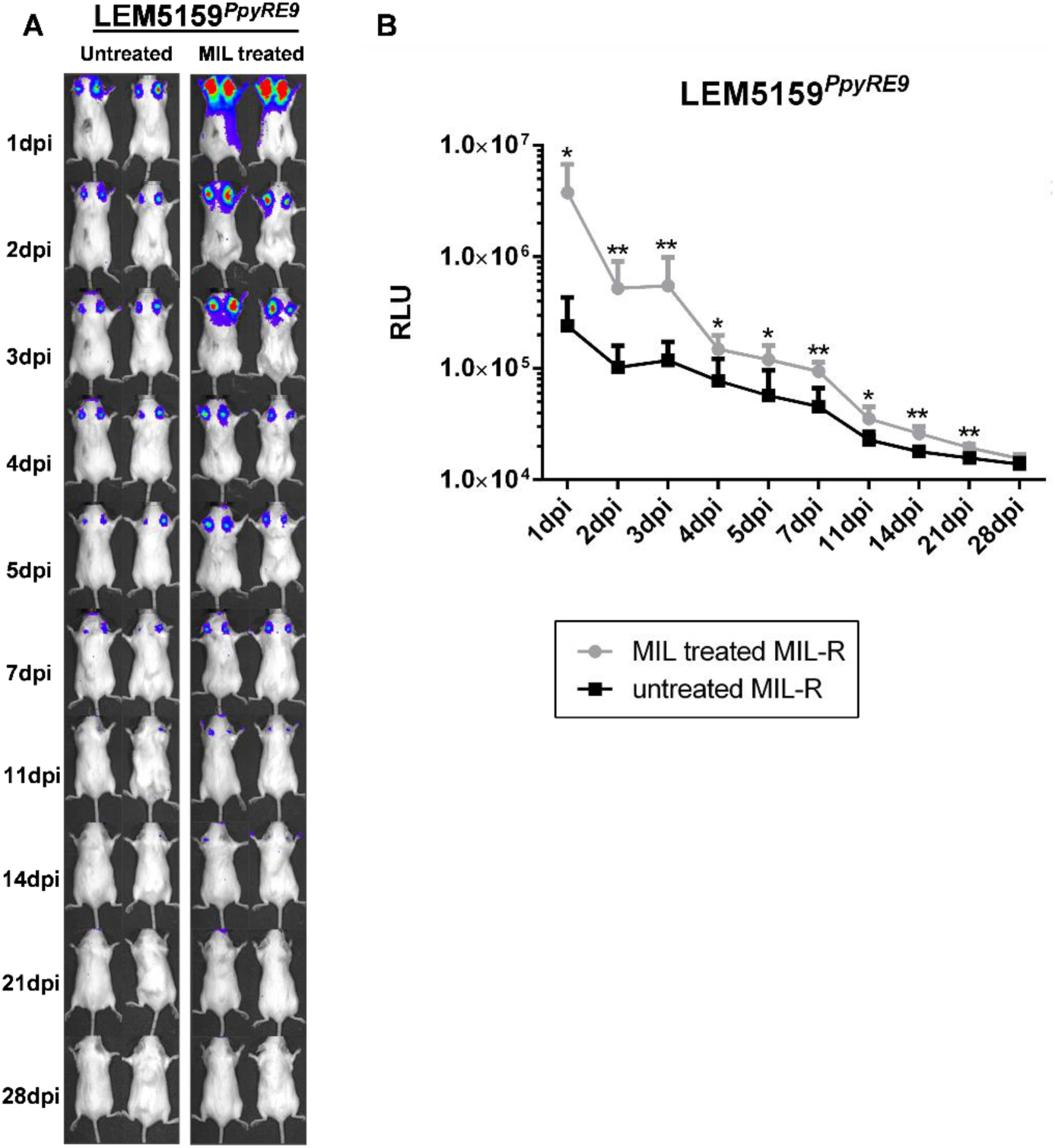
Effect of *in vivo* MIL treatment on intradermal infection with MIL-R parasites. (A) Bioluminescent imaging using an exposure time of 10 min of LEM5159*^PpyRE^*^9^ i.d. infection in MIL treated and untreated BALB/c mice. Mice were treated with 40 mg/kg MIL via oral gavage for 5 consecutive days starting one day before infection. (B) Mean RLU values at the inoculation site during the first 4 weeks of LEM5159*^PpyRE9^* i.d. infection in MIL treated and untreated BALB/c mice. Each infection group consisted of 3 BALB/c mice. Results are expressed as mean ± SD (* p≤0.05, ** p ≤0.01).

## Discussion

Miltefosine (MIL) is currently the only oral drug available to treat VL with initial cure rates above 94% upon introduction in the Indian subcontinent (24, 51). Within one decade of use treatment failure rates during a 12 month follow-up increased to 10% in India and up to 20% in Nepal (32, 33). Despite this evolution, only a limited number of fully resistant field isolates have been described, of which two MIL-R *L. infantum* isolated from French HIV-infected VL patients who received several successive treatments with MIL and liposomal amphotericin B (35,36,39,52). This might suggest a connection between the emergence of MIL-resistance in *L. infantum* parasites and the immunocompromised state of the host. Furthermore, the unresponsiveness of Brazilian *L. infantum* strains to MIL was recently attributed to the absence of a MIL-sensitivity locus that was shown to be present in the genomes of all sequenced *L. infantum* and *L. donovani* isolated in the Old World (53). The scarce number of resistant field isolates is surprising considering that full resistance can be obtained via single mutations in a aminophospholipid transporter complex, encoded by the *MT/Ros3* genes (37, 40). It was shown that experimentally induced MIL-R *L. major* and *L. infantum* are hampered by a severe *in vivo* fitness loss which renders them unable to mount a successful infection in mice (40, 42). Given this limited information, the present study investigated the impact of acquired resistance on *in vivo* host cell infection and initiation of immune responses using the experimentally induced MIL-R *L. infantum* LEM3323 line carrying an inactivating 2 bp deletion in the *LiMT* gene (37).

By generating fluorescent/bioluminescent reporter strains, the *in vivo* fitness loss of MIL-R parasites in BALB/c and C57Bl/6 mice was confirmed by a rapidly decreasing liver signal and the absence of successful dissemination to the spleen and bone marrow. When exploring the host cell infectivity, a higher number of infected cells was detected at 1 dpi in the liver and spleen of MIL-R infected mice. The number MIL-R infected cells progressively decreased by 21 dpi whereas the number of MIL-S infected cells increased as shown by tissue imprints and bioluminescent imaging corresponding to parasite expansion in the liver. Flow cytometric detection of the DsRed reporter suggests the occurrence of a progressively quiescent state and decreased reporter gene expression over the course of infection. Flow cytometry analysis of different infected cell populations further revealed differences in cellular tropism between MIL-S and MIL-R which were correlated with an increased influx and concomitant infection of monocytes, neutrophils and dendritic cells by MIL-R. As a result, MIL-R less efficiently established in F4/80^+^ macrophages.

Compared to MIL-S, MIL-R parasites also induced elevated IFN-γ levels at 1 dpi which was attributed to an increased activation of hepatic NK and NKT cells. This activation contributed to the reduced survival of MIL-R parasites *in vivo*. Both NK and NKT cells have previously been shown to produce IFN-γ during VL infection onset in mice: NKT cells are activated in a CD1d-dependent manner by parasite lipid antigens, such as LPG and glycosphingolipid (GSL) presented on dendritic cells and Kupffer cells (21,54–56). Moreover, GSL-induced relocation of CD1d molecules into lipid rafts facilitates the interaction with NKT cells (54). NK cells are known to become activated by LPG through engagement of this phosphoglycan with TLR2 (57). Mutations in the *LiMT/LiRos3* transporter disrupts the transportation of lipids affecting the lipid asymmetry of the plasma membrane by exposing phosphatidylethanolamine and phosphatidylcholine on the outer membrane (44). MIL-R parasites were shown to contain more saturated phospholipid acyl chains and more short alkyl fatty acids, resulting in a higher membrane rigidity (58, 59). We hypothesize that this altered lipid composition of the MIL-R membrane directly results in increased NK and NKT cell activation and enhanced presentation of lipid antigens by CD1d molecules on dendritic cells and Kupffer cells. However, despite the increased survival of MIL-R parasites in the absence of activated NK and NKT cells, fitness was not fully restored indicating the contribution of additional factors to the decreased *in vivo* survival. Our results also indicate that neutrophils contribute to the early IFN-γ burst against the MIL-R strain. Several other studies already suggested the involvement of neutrophils in regulating the IFN-γ production during VL infection (60–62) although the exact mechanisms remain to be elucidated. Neutrophils are known to influence NKT cell activation. In neutrophilic mice, the number of hepatic NKT cells was shown to be reduced as well as the proportion of IFN-γ-producing NKT cells in response to αGalCer (63). This reduced IFN-γ production was correlated with a reduced *T-bet* expression in NKT cells. Neutrophils can also indirectly influence IFN-γ production by NKT cells. In the skin, *L. major* infected neutrophils were shown to be efficiently captured by dendritic cells (64). Similar interactions have not yet been addressed during VL infection but may contribute to increased CD1d presentation capacity of dendritic cells resulting in an increased NKT cell activation. The observation of reduced parasite numbers in the absence neutrophils corroborates the concept of a dual role for neutrophils during MIL-R infection. Neutrophils were shown to be the major infected population during the first 24 hours of intradermal inoculation with *L. infantum* and are regarded as an important transient safe haven that facilitates passage of parasites to macrophages (65–67). However, they are also known to contribute to protective antileishmanial immune responses as evidenced by a delayed hepatic granuloma formation, decreased expression of iNOS and IFN-γ and increased production of IL-4 and IL-10 in the absence of neutrophils (61,66–69).

Our lab recently described drug-dependency of the MIL-R parasite whereby the *in vivo* fitness loss in BALB/c mice was partially restored by exposure to MIL (42). Analogous experiments in C57Bl/6 mice confirmed this drug effect. The present results showed that MIL-exposure lowers NK and NKT cell activation resulting in a decreased IFN-γ production early in the infection. The exact mechanism behind this MIL-induced modulation remains unclear, although interactions of MIL with the parasite’s plasma membrane are likely to play a role. The interaction and incorporation of MIL in lipid monolayers have been described (70, 71) and it was shown that MIL can increase the lipid dynamics of *L. amazonensis* membranes (72). MIL is known as a lipid raft modulating agent (73) where the condensed microdomains of lipid rafts are suggested to incorporate high quantities of MIL (58, 59). This incorporation may affect the presentation of parasite antigens. The effect of MIL on other antiparasitic mechanisms should also be considered. For example, a recent study with MIL-tolerant *L. donovani* parasites from relapse patients showed a decreased production of ROS by infected macrophages following MIL exposure (34). The drug-dependency of MIL-R parasites was further confirmed using the natural MIL-R LEM5159 strain via an intradermal infection model which better represents the natural route of infection. In this model with the natural MIL-R parasite, MIL treatment also resulted in higher dermal parasite burdens.

Altogether, these observations show that MIL-R parasites induce an enhanced antileishmanial innate immune response. Together with a different cellular tropism and the inability to successfully survive and multiply inside macrophages, this results in the rapid clearance of MIL-R from the liver and spleen. This increased immunogenicity might account for the scarce number of MIL-R field isolates and possibly explains why the few known *L. infantum* MIL-R field isolates were isolated from HIV-infected VL patients. Furthermore, these observations emphasize the risk of MIL treatment in sustaining infections with MIL-R parasites and provide an immunological basis for the attenuated and drug-dependent phenotype of resistant parasites.

## Material and methods

### Ethics statement

The use of laboratory animals was carried out in strict accordance to all mandatory guidelines (EU directive 2010/63/EU on the protection of animals used for scientific purpose and the declaration of Helsinki) and was approved by the Ethical Committee of the University of Antwerp (UA-ECD 2015-90 and 2017-04).

### Animals

Female Swiss, BALB/c and C57Bl/6 mice (6-8 weeks old) were purchased from Janvier Laboratories (Le Genest-Saint-Isle, France). C57Bl/6 IFN-γ reporter mice with endogenous polyA transcript (GREAT) and were obtained from Prof. C. De Trez (VUB, Belgium). Mice were randomly allocated and housed in individually ventilated cages with laboratory rodent food (Carfil, Belgium) and drinking water *ad libitum*.

### Parasites and transfections

The clinically MIL-resistant *L. infantum* strain LEM5159 (MHOM/FR/2005/LEM5159) (36, 37), the MIL-susceptible *L. infantum* strain LEM3323 MIL-S (MHOM/FR/96/LEM3323) and its MIL-resistant isogeneic counterpart LEM3323 MIL-R were previously rendered bioluminescent by transfection with the *PpyRE9* gene (42). Both strains were additionally transfected with the *DsRed* gene that was cloned from the pssuINTDsRed vector which was obtained through Prof. C. De Trez (VUB, Belgium). A homologous overlap for Gibson assembly in the pLEXSY-Sat2.1 vector (Jena Bioscience, Germany) was added to the *DsRed* gene during cloning using the primers FP 5’-CGC TCC TGC TTT CCT TGC TGT GCC TTG CCA CCA GAT CTG CAT GGC CTC CTC CGA GG-3’ and RP 5’-AGA AGG CGA CGT GAA AGG AAC AAG AAA GGA GGA GGA GGG CCT ACA GGA ACA GGT GGT GG-3’. pLEXSY-Sat2.1-DsRed was digested using SwaI restriction enzyme and the 2.8 kb fragment was gel-purified using the QIAquick Gel Extraction Kit (Qiagen, Germany) followed by ethanol precipitation and resuspension in cytomix transfection buffer (120 mM KCl; 0.15 mM CaCl_2_; 10mM KH_2_PO_4_; 25mM HEPES; 2mM EDTA; 5mM MgCl_2_; pH 7.6) at 1 µg/µL. For each strain, 10 µg of the linearized construct was used for the electroporation of 1×10^8^ procyclic promastigotes (twice at 25 µF and 1500V with 10s interval) using the Bio-Rad GenePulse Xcell electroporation unit (Bio-rad Laboratories, USA). MIL-S*^PpyRE9/DsRed^* and MIL-R*^PpyRE9/DsRed^* transfectants were selected under Nourseothricin (Jena Bioscience, Germany) pressure and subcultured twice weekly in HOMEM promastigote medium.

### Animal infections

Mice were infected with MIL-S*^PpyRE9/DsRed^* or MIL-R*^PpyRE9/DsRed^* double reporter parasites by intravenous (i.v.) injection of 1×10^8^ metacyclic promastigotes. In some experiments, MIL-R*^PpyRE9/DsRed^* parasites were preconditioned for 5 days with 40 µM MIL prior to infection. To assess the viability of stationary phase, metacyclic MIL-S and MIL-R parasites prior to infection, parasites were stained with propidium iodide (Thermo Fisher scientific, USA) and analyzed on a MACSQuant 10 Analyzer (Miltenyi Biotec, Germany) and using FlowLogic software (Miltenyi Biotec, Germany). As a positive control, heat-killed (15 min. at 65°C) were included. Both MIL-S*^PpyRE9/DsRed^* and MIL-R*^PpyRE9/DsRed^* stationary phase cultures had similar percentages of dead parasites (1.097 ± 0.294% and 0.971 ± 0.591%, respectively) (Fig. S1) indicating that any differences observed during the *in vitro* and *in vivo* infection could not be attributed to differences in inoculum viability. Parasite burdens were monitored *in vivo* using bioluminescent imaging (BLI) of mice with the IVIS System (PerkinElmer, Belgium) throughout early infection. In some experiments, 1×10^7^ metacyclic promastigotes were inoculated intradermally (i.d.) in the ear of BALB/c mice that were treated for 5 consecutive days, starting 1 day before infection, with 40 mg/kg of MIL via oral gavage. Mice were imaged for 15 minutes starting 3 minutes after i.p. injection of 150 mg/kg D-luciferin (Beetle Luciferin Potassium Salt, Promega, USA). Images were analyzed using LivingImage v4.3.1 software (PerkinElmer, Belgium) by quantifying the signal as relative luminescence units (RLU) within regions of interest corresponding to the ear, liver, spleen and bone marrow. At each time point, blood was collected via the tail vein and plasma was stored at -80°C until further analysis.

### Cell isolation

Infected C57Bl/6 and/or C57Bl/6 IFN-γ reporter mice were sacrificed at 1, 7 and 21 dpi. Spleen and liver were isolated after transcardial perfusion with 10 mL KREBS Henseleit solution supplemented with 10 U/mL heparin (Sigma-Aldrich, USA) (74). Tissue imprints of infected livers and spleens were made on glass slides, fixed with methanol, Giemsa-stained and evaluated microscopically to determine the level of infection. Livers were homogenized using a liver dissociation kit (Miltenyi Biotec, Germany) according to the manufacturer’s instructions. Briefly, livers were mechanically disrupted in 5 mL DMEM medium (Thermo Fisher scientific, USA) containing liver dissociation enzymes, using the gentleMACS Dissociator (Miltenyi Biotec, Germany) and incubated at 37°C for 30 min. The digested tissue was homogenized and filtered through a 100 µM filter. Spleens were dissociated in 10 mL FACS medium (PBS, 5% FCS, 2mM EDTA, pH 7.2) by mechanical disruption. Subsequently, spleen cells were centrifuged (10 min, 400×*g*, 4°C) and the pellet was treated with ACK lysis buffer (0.15 M NH_4_Cl, 1 mM KHCO_3_, 0.1 mM Na_2_-EDTA). All cell suspensions (spleen and liver) were centrifuged (10 min, 400×*g*, 4°C) and the pellet was resuspended in FACS medium at a concentration of 2×10^7^ cells/ml for flow cytometry analysis.

### Flow cytometry

Spleen and liver cells were incubated (20 min, 4°C) with anti-Fc-gamma blocking antibody (clone 2.4G2) (BD Biosciences, Belgium) and were subsequently stained with fluorescent conjugated antibodies (30 min, 4°C, in the dark): Ly-6G FITC (clone 1A8), F4/80 PE-Cy7 (clone BM8), CD11b Pacific Blue (clone M1/70), CD11c PerCP-Vio700 (clone N418), Ly-6C APC (clone AL-21), CD45 APC-Cy7 (clone 30-F11), CD4 FITC (clone GK1.5), CD45 PerCP (clone 30F11), CD8 PE-Cy7 (clone 53-6.7), TcRβ APC (clone H57-597), NK1.1 APC-Cy7 (clone PK136) and BV421-conjugated, PBS-57 loaded, mouse CD1d tetramer. Antibodies were purchased from either BD Biosciences, BioLegend (USA) or Miltenyi Biotec and mouse CD1d tetramers were obtained from the Tetramer Core Facility (NIH/NIAID, USA). Viability was assessed using LIVE/DEAD Fixable Aqua Dead Cell Stain (Thermo Fisher scientific, USA). Cells were subsequently washed and measured on a MACSQuant 10 Analyzer (Miltenyi Biotec, Germany) and results were analyzed using FlowLogic software (Miltenyi Biotec, Germany). Different liver and spleen cell populations were identified using the gating strategy described in Fig. S2. Total cell numbers were calculated using volumetric measurement of living CD45^+^ cells for each sample.

At several time points over the course of infection, different DsRed^+^ cell populations were sorted using a BD FACS Melody apparatus (BD Biosciences, Belgium). Sorted cells were centrifuged for 10 minutes at 1,000×g and pelleted on a glass slide using a cytospin 4 cytocentrifuge (Thermo Fisher scientific, USA). Glass slides were air dried, subjected to a 5-minute fixation in methanol and staining with Giemsa solution for 15 minutes and analyzed microscopically.

### Cell depletion and neutralization experiments

Depletion and neutralization experiments were based on previously established protocols in mice. Neutralization of the early IFN-γ burst was done by a single high dose (1 mg) injection of an anti-IFN-γ neutralizing rat-anti mouse monoclonal antibody (clone R4-6A2; BioXCell, USA) or an IgG1 isotype control antibody (clone HRPN; BioXCell, USA) one day prior to infection (75, 76). In C57Bl/6 mice, NK and NKT cells were depleted using anti-NK1.1 depleting rat anti-mouse monoclonal antibody (clone PK136; BioXCell, USA). Hundred µg of the antibody or an IgG2a isotype control antibody (clone C1.18.4; BioXCell, USA) was given intraperitoneally (i.p.) on days -7, -4, -1, 1, 4 and 7 (77, 78). For neutralization of CD1d presentation to NKT cells, BALB/c and C57Bl/6 mice were treated i.p. with 75 µg neutralizing anti-mouse CD1d antibody (clone 19G11; BioXCell, USA) or an IgG1 isotype control (clone HRPN; BioXCell, USA) on days -7, -4, -1, 1, 4 and 7 (79). For the depletion of neutrophils, BALB/c mice received 75 µg of depleting anti-Ly-6G antibody (clone 1A8) (BioXCell, USA) or isotype IgG2a isotype control antibody (clone 2A3; BioXCell, USA) i.p. on days -4, -1, 1, 4 and 7 (80, 81).

### Serum analysis

Serum levels of IFN-γ, IL-10 and TNF-α were determined using the V-PLEX Custom Mouse Cytokine assay (Meso Scale Discovery, Rockville, USA) according to the manufacturer’s instructions.

### RNA extraction

Dissected liver and spleen samples were treated with RNALater (Qiagen, Germany) according to the manufacturer’s instructions and stored at -80°C. Tissue samples were thawed and RNA extraction was performed using the Qiagen RNA extraction kit (Qiagen, Germany). Briefly, samples were homogenized in 100 µL RLT buffer using a pestle rotor. After homogenization, 500 µL RLT buffer was added and samples were centrifuged for 3 minutes at maximum speed after which the supernatant was loaded on a gDNA Eliminator spin column and centrifuged for 30 seconds at 10,000×g. 1 volume of 70% ethanol (50% for liver) was added to the flow through and loaded onto a RNeasy spin column followed by centrifugation for 15 seconds at 10,000×g. The flowthrough was discarded and columns were washed with RW1 and RPE buffer. RNA was eluted in RNA-free water following a centrifugation for 1 minute at 10,000×g. Concentration of RNA samples was measured by absorbance at 260 nm using a NanoDrop spectrophotometer (Isogen Life Science, The Netherlands). Absorbance ratios at 260/280 nm and 260/230 nm were determined as a measure for RNA purity and integrity. Samples were stored at -80°C until qPCR analysis.

### Real-time quantitative PCR

For real-time quantitative PCR (RT-qPCR) detection of IFN-γ, IL-10, TNF-α and IL-1β encoding transcripts in liver and spleen, validated primers were obtained from an online repository (http://www.rtprimerdb.org) (table 1). 1:10 dilutions of RNA samples were analyzed using SensiFast SYBR Hi-ROX One-Step kit (Bioline, UK) and 400 nM primer concentrations. PCR conditions comprised an initial polymerase activation at 96°C followed by 40 cycles, each consisting of a denaturation step at 95°C for 15 seconds, 30 seconds annealing at 60°C and 30 seconds elongation at 72°C. The specificity of the primers was assessed by verifying the presence of a single amplicon by melting curve analysis. *Eukaryotic translation elongation factor 2* (*eef2*)(82) was used as reference gene for normalization in the Q base Plus Software (Biogazelle, Belgium). The relative expression levels were analyzed using Graphpad Prism software v7 (GraphPad, USA).

**Table 1:**
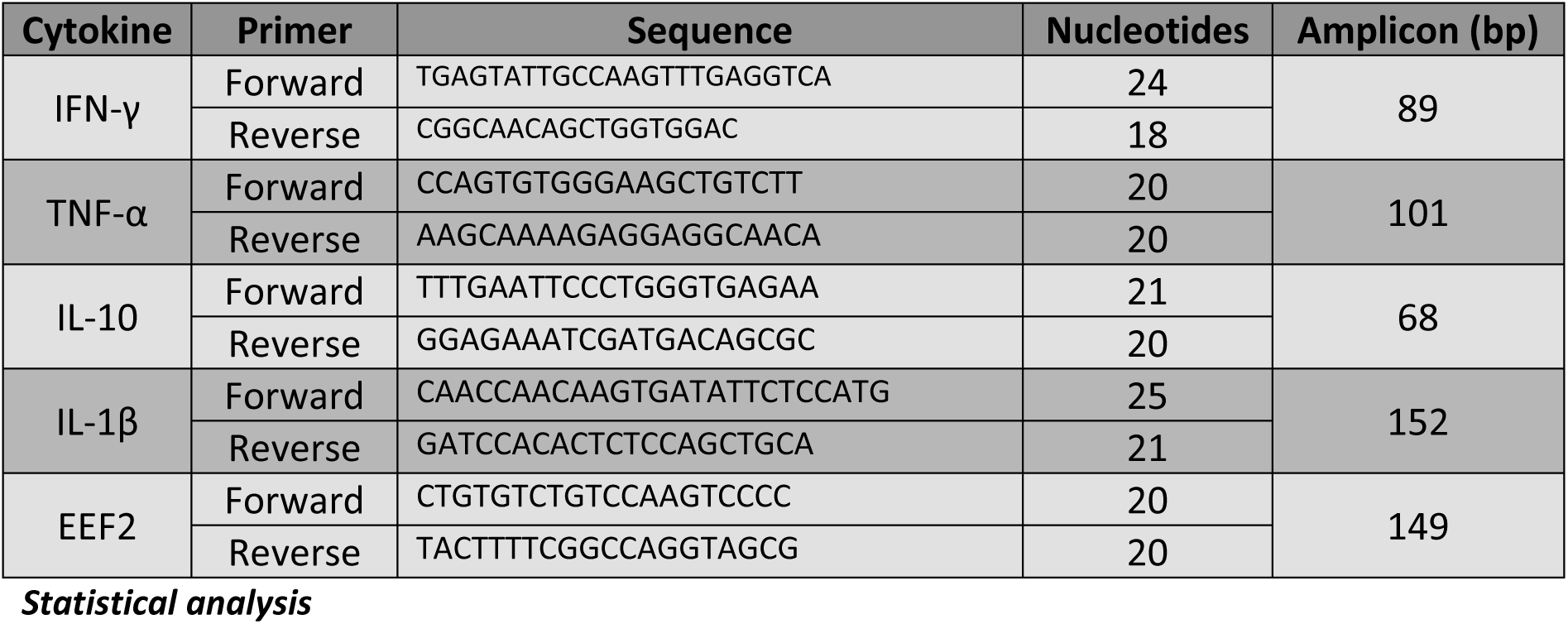
Sequence of RT qPCR primers for the detection of several inflammatory genes.

Statistical analysis was performed using the Mann-Whitney U test in IBM SPSS Statistics v25 (IBM, Armonk, USA). Values are expressed as mean ± standard deviation (SD) and values of p ≤0.05 are considered statistically significant.

## Acknowledgements

Following reagents were obtained via the NIH Tetramer Core Facility: BV421 and APC conjugated, PBS57 loaded CD1d tetramers. This work was funded by the Fonds Wetenschappelijk Onderzoek Vlaanderen [FWO No. 1136417N (L.V.B.), 12I0317N (S.H.)], Research Funds of the University of Antwerp [TT-ZAPBOF 33049 (G.C.) and TOP-BOF 35017 (LM, GC)]. The authors thank Laurence Lachaud (Laboratoire de Parasitologie-Mycologie et Centre National de Référence des Leishmanioses, Montpellier, France) for providing the LEM3323 clinical isolate. We acknowledge Prof. Carl De Trez (VUB, Brussels, Belgium) for providing the IFN-γ reporter mice. LMPH is a partner of the Excellence Centre ‘Infla-Med’ (http://www.uantwerpen.be/infla-med).

